# Unilateral Perforant Path Transection Does Not Alter Lateral Entorhinal Cortical or Hippocampal CA3 *Arc* Expression

**DOI:** 10.1101/2022.04.14.488420

**Authors:** Tara L. Cooper, John J. Thompson, Sean M. Turner, Cory Watson, Katelyn N. Lubke, Carly N. Logan, Andrew P. Maurer, Sara N. Burke

## Abstract

It is well established that degradation of perforant path fibers is associated with age-related cognitive dysfunction and CA3 hyperactivity. Whether this fiber loss triggers a cascade of other functional changes within the hippocampus circuit has not been causatively established, however. Thus, the current study evaluated the effect of perforant path fiber loss on neuronal activity in CA3 and layer II of the lateral entorhinal cortex (LEC) in relation to mnemonic similarity task performance. Expression of the immediate early gene *Arc* was quantified in rats that received a unilateral right hemisphere transection of the perforant path or sham surgery that cut the cortex but left the fibers intact. Behavior-related expression of *Arc* mRNA was measured to test the hypothesis that fiber loss leads to elevated activation of CA3 and LEC neurons, as previously observed in aged rats that were impaired on the mnemonic similarity task. Transection of perforant path fibers, which has previously been shown to lead to a decline in mnemonic similarity task performance, did not alter *Arc* expression. *Arc* expression in CA3, however, was correlated with task performance on the more difficult discrimination trials across both surgical groups. These observations further support a link between CA3 activity and mnemonic similarity task performance but suggest the reduced input from the entorhinal cortex to the hippocampus, as observed in old age, does not causatively elevate CA3 activity.

## 1 INTRODUCTION

The perforant path is the primary fiber projection from the cortex to the hippocampus. A major component of this cortical to hippocampal white matter tract are the axons from layer II entorhinal cortical neurons that synapse in CA3 and the dentate gyrus (Amaral and Witter, 1995). This fiber projection is known to degrade in advanced age. This was first established by the observation that presynaptic fiber potential for a given stimulation intensity of perforant path axons was reduced in aged compared to young rats (Barnes and McNaughton, 1980; Foster et al., 1991). Later, stereological electron microscopy was used to show that in aged compared to young rats there were fewer synapses in the stratum lacunomsum of CA3 (Adams et al., 2010) and the dentate gyrus molecular layer (Geinisman et al., 1992). These dendritic regions of CA3 and dentate gyrus both receive direct input from layer II neurons in the entorhinal cortex. Age-related reductions in perforant path white matter integrity have also been shown in studies using magnetic resonance imaging in human study participants (Stoub et al., 2005; Yassa et al., 2010; Yassa et al., 2011a; Rogalski et al., 2012(Yassa et al., 2011a; Bennett et al., 2015; Bennett and Stark, 2016).

While the number of neurons in the hippocampus (Rapp and Gallagher, 1996) and entorhinal cortex (Rapp et al., 2002) does not change with age, age-related deficits have been associated with hyperactivity in the CA3 subregion in rodents (Wilson et al., 2005; Robitsek et al., 2015; Lee et al., 2021a), and nonhuman primates (Thomé et al., 2016). Additionally, fMRI studies in human study participants have reported that the BOLD signal in CA3/dentate gyrus is higher in older compared to younger adults (Yassa et al., 2010; Yassa et al., 2011a; Reagh et al., 2018). This aberrant activity is hypothesized to underlie poor cognition and is exacerbated in persons with Mild Cognitive Impairment and Alzheimer’s Disease (Bakker et al., 2012; Bakker et al., 2015; Tran et al., 2017). While correlations between CA3 hyperactivity and poor cognitive outcomes in old age have been observed across species, the cellular changes that provoke this increased activity in aged animals is unknown.

One hypothesis is that perforant path fiber disruptions, discussed above, cause a reduction in the recruitment of interneuron activity and lead to an imbalance between inhibition and excitation. In support of this idea, CA3 hyperactivity correlates with the magnitude of perforant path degradation in humans (Yassa et al., 2011). These data suggest that disruptions in this white matter tract could initiate a cascade of events that leads to CA3 hyperactivity and cognitive dysfunction. In support of this idea, aged rats that performed poorly on a rodent analog of the mnemonic similarity task (Stark et al., 2013; Johnson et al., 2017), showed a higher proportion of cells that were positive for mRNA of the immediate-early gene *Arc*, a neuronal marker for plasticity, in both CA3 and in the lateral entorhinal cortical (LEC) neurons that project to the hippocampus (Maurer et al., 2017). This observation suggests that a common underlying mechanism could be related to both LEC and CA3 dysfunction in cognitively impaired aged animals. Whether this is due to perforant path fiber disruption, however, has not been empirically examined.

Aged rodents (Burke et al., 2010; Johnson et al., 2017), monkeys (Burke et al., 2011; Johnson et al., 2017) and humans (Stark et al., 2013; Stark et al., 2015; Stark and Stark, 2017) are less accurate at the discrimination of similar stimuli that share features compared to young animals. Specifically, older study participants are more likely to classify novel lures as familiar when they share characteristics with a stimulus that has previously been experienced. This behavioral deficit can be recapitulated in young animals following a unilateral right hemisphere perforant path transection (Burke et al., 2018). This finding suggests that degradation of perforant path fibers in young animals may model cognitive deficits seen in aging that result in impaired behavior and discrimination ability. Potential changes in neuronal activity underlying behavioral deficits seen in young animals following disconnection of perforant path fibers and how these changes compare to data in aged animals showing CA3 hyperactivity have not been examined, however. Thus, the current study interrogated whether loss of perforant path fibers recapitulates CA3 hyperactivity seen in aged animals by quantifying the expression of the immediate-early gene *Arc* in relation to performance on the rodent mnemonic similarity task in rats that had received a unilateral perforant path transection.

In the current study, rats that received a right hemisphere perforant path transection and control animals (sham surgery that cut the cortex, leaving fibers intact) performed two epochs of the rodent mnemonic similarity task. In the different epochs, the amount of feature overlap of the two objects to be discriminated was either low (70%) or high (90%), and the order of epochs was counterbalanced across animals. *Arc* expression in relation to behavior was then examined in CA3 and cortical layer II of the LEC. The subcellular localization of the immediate-early gene *Arc* can be used to determine which neural ensembles across the brain were active during 2 distinct episodes of behavior. *Arc* is first transcribed within the nucleus of neurons 1 to 2 minutes after patterned synaptic activity associated with neuronal plasticity and cell firing. *Arc* mRNA then translocates to the cytoplasm approximately 15 to 20 minutes after cell firing, which allows for cellular activity during 2 epochs of behavior, separated by a 20-minute rest to be represented within a single neural population (Guzowski et al., 1999). An observed increase in CA3 *Arc* expression following fiber transection would suggest a causative relationship between white matter damage and CA3 hyperactivity. Contrary to this hypothesis, perforant path fiber damage did not impact CA3 or LEC *Arc* expression suggesting that compensatory mechanisms were employed to normalize circuit activity in the face of fiber damage.

## 2 MATERIALS AND METHODS

### 2.1 Animals

Experiments used 37 young (4-6 months old) male and female F344 x Brown Norway F1 hybrid rats (17 male/20 female) that were 4 months old at arrival to the UF vivarium. Rats were single-housed on a reversed 12-hour light/dark cycle and experiments were conducted during the dark phase. When the rats arrived at the facility, they were given 7 days to acclimate prior to initiating food restriction. All rats were then handled for an additional 7 days prior to behavioral shaping. The behavioral training, testing procedures and results are as described in (Burke et al., 2018). Briefly, experimentation and training occurred at least 5 days per week at approximately the same time each day. Before experimentation, rats were placed on a restricted diet of a moist chow (20±5g; ∼39kcal/day; Teklad LM-485, Harlan Labs) to provide a food-based motivation for behavioral experiments. Drinking water was given *ad libitum*. Once rats had reached 85% of their baseline body weight, they began habituation to the testing procedures. All animals handling procedures and protocols were carried out in accordance with NIH *Guide for the Care and Use of Laboratory Animals* and approved by the Institutional Animal Care and Use Committee at the University of Florida.

### 2.2 Apparatus

All experimentation and training procedures were conducted in an L-shaped track as shown in **Figure 1A** and previously described in (Johnson et al., 2017). The track contained a starting area, narrow track measuring 84 cm x 10.2 cm with 6.4 cm walls on both sides, and a choice platform measuring 32 cm x 24 cm also with walls 6.4 cm in height. There were two food reward wells, each 2.5 cm in diameter, centered in the choice platform 12 cm apart, 7.6 cm from the back wall, and recessed 1 cm into the floor. All experiments took place in a dimly lit testing room with a red 60W light bulb near the starting area and another shining on the choice platform. An additional white 60W light bulb was placed in a lamp that aimed at the ceiling in the corner of the room to facilitate video monitoring and as an additional light source for illumination of the object stimuli used in the experiments and in previous experiments (Burke et al., 2018). A webcam (Logitech; Newark, CA, United States) was mounted 50cm above the choice platform to produce video recordings of discrimination trials. Trials were recorded using custom software (Collector; Burke/Maurer, Gainesville, FL, United States). Additionally, a white noise machine was used during all testing session, in order to reduce the likelihood that external noise would cause distraction.

**Figure 1.**
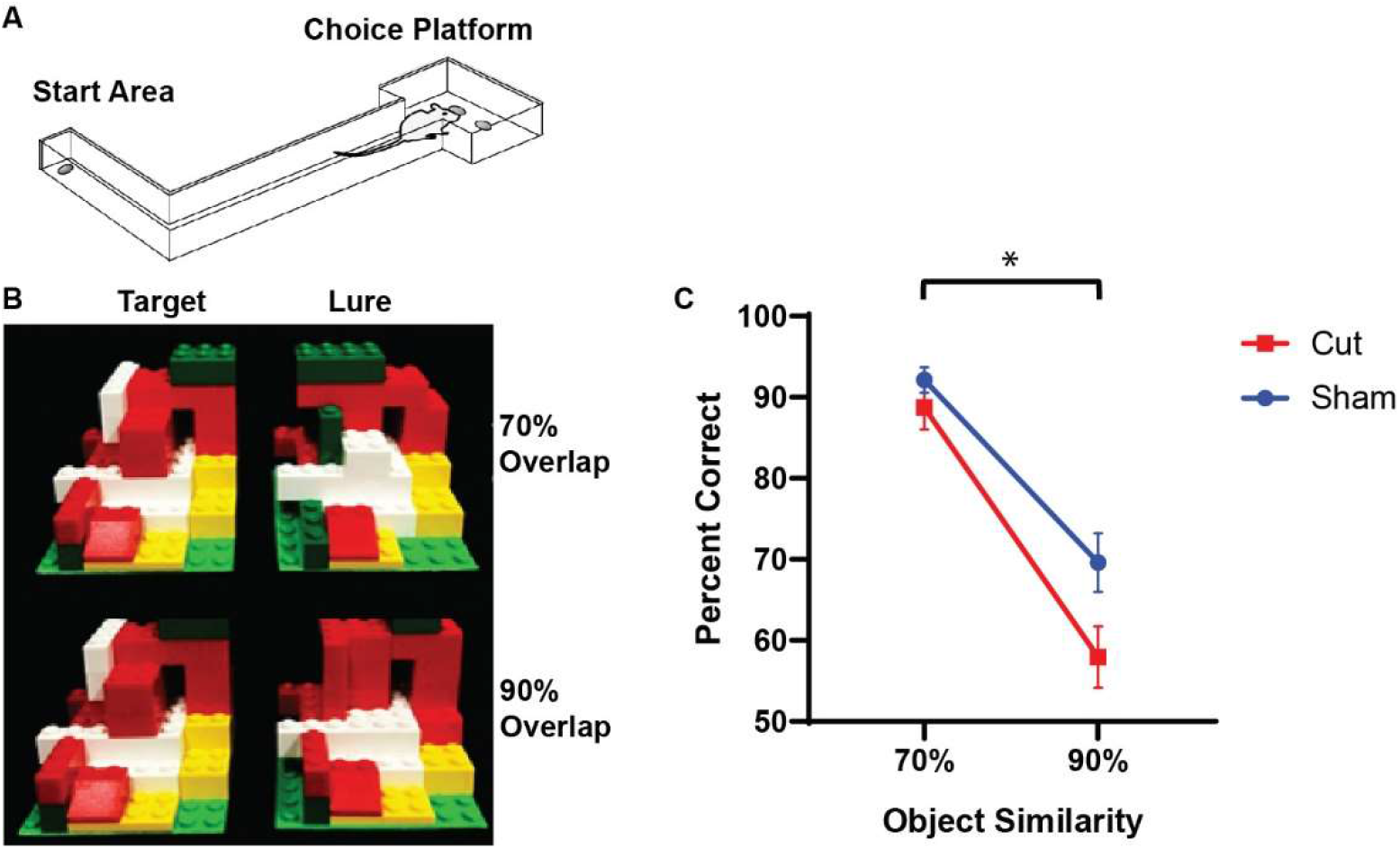
The rodent mnemonic discrimination task. (**A**) Schematic of the L-shaped apparatus used for both training and testing discrimination ability in rats. (**B**) The LEGO block object pairs used in testing. Rats were trained to identify the target object as correct compared to either a 70% (top) or 90% (bottom) visual overlap object. (**C**) Mean percent correct trials on target-lure discrimination task. Rats performed worse on the higher similarity lure trials (F[1,28] = 62.361, p < 0.001). Furthermore, there was a main effect of experimental surgical condition (F[1,28] = 6.072, p = 0.020), with Sham animals having more correct trials compared to the Cut rats. There was no main effect of sex on performance (F[1,28] = 0.752, p = 0.393). Error bars are ± one standard error of the mean (SEM). * Indicates significant effect.

### 2.3 Habituation and Shaping

Prior to the beginning of habituation, rats were given Froot Loops cereal pieces along with the rat chow, in order to familiarize the animals with the reward. At the beginning of habituation, rats were placed inside the L-shaped track near the starting area and explored the track for 10 minutes, during which they retrieved Froot Loops cereal pieces throughout the track (Kellogg’s; Battle Creek, MI, United States). Habituation usually lasted anywhere from 1-3 days and shaping began when rats could explore the complete L-shaped track for rewards without external assistance. During shaping, rats learned to alternate between the starting area and the choice platform. This was accomplished by placing Froot Loops in the choice wells, placing the animals at the start area, and allowing them to alternate between the two for reward. This process was repeated until a rat could achieve 32 alterations in 20 min (Burke et al., 2018).

### 2.4 Standard Object Discrimination and LEGO® Block Object Discrimination

In the first discrimination task, all rats were given the option of choosing between two distinct objects (Bunny-Rooster) that had variations in size, shape, color, and texture. Half of the rats were rewarded for selecting the bunny and the other half for the rooster. Similar to the previous stages of training, rats were placed in the starting area, ran down the track, and would displace one of the objects, where the correct object and corresponding food well were randomly alternated during the course of the experiment. Upon a correct choice between the objects, the rats were allowed access to the food reward both at the food well and at the starting area, so as to start the next trial, whereas incorrect responses resulted in removal of the Froot Loops piece and no food reward in the well (Burke et al., 2018). This session continued until rats completed 32 trials, and this stage of testing occurred daily until rats reached criterion performance of choosing the correct object 26 out of 32 trials (81.25% correct). To prevent rats from choosing an object solely based on the scent of the object, each object was cleaned thoroughly with 70% ethanol and was rotated out with other exact copies.

Once rats had reached criterion on the standard task, they were then moved onto the LEGO block object Discrimination Task. In this task, rats were given objects constructed out of LEGO blocks and had to discriminate between LEGO objects with similar volume and size (see Figure 1B). The front face of the LEGO objects, that which was visible to rats in the testing space, varied in composition. Methods used to train the rats to discriminate between the two LEGO objects were identical to the methods for classic object discrimination. Training continued until a rat was able to reach the criteria of 81.25% correct (26 correct of 32 trials). Once they had reached criterion performance, animals were trained to discriminate a different novel LEGO object pair in which the correct object was the target for the LEGO block Mnemonic Discrimination task. Once the animals had reached criterion performance of 26 correct of 32 trials, for a final time, rats underwent surgical transection of the perforant path.

### 2.5 Perforant Path Transection & Cholera Toxin Subunit B Injection

Each rat underwent stereotaxic surgery under isoflurane anesthesia (1–3%) to either unilaterally cut the perforant path fibers in the right hemisphere or a sham surgery in which only right hemisphere cortex was cut. All animals were injected with the retrograde tracer Cholera Toxin Subunit B in both the left (Alexa Fluor 647 Conjugate, C34778, Invitrogen, Carlsbad, CA, United States) and right (Alexa Fluor 488 Conjugate, catalog #: C22841, Invitrogen, Carlsbad, CA, United States) hippocampus. Cholera Toxin Subunit B conjugated with Alexa Fluor 488 was injected into the right hemisphere to verify the knife cut in animals that received the transection and intact fibers in sham animals. CTB with the Alexa Fluor 647 conjugate was used as a positive control to ensure that the tracer was visible in the entorhinal cortex of the non-lesioned hemisphere. During surgery, an incision was made to expose Bregma and Lambda. A 2.5 mm wide craniotomy was drilled in the skull centered at 1.7 mm anterior to the intra-aural line in males and extending laterally in the right hemisphere from 0.5 to 5.0 mm lateral to the midline. For females, to adjust for differences in skull and brain size, coordinates were modified based on the distance between bregma and the intra-aural line (Paxinos et al., 1985). In female rats, the craniotomy was centered at 1.9 mm anterior to the intra-aural line and extending from 0.5 to 4.75 mm lateral to the midline. Knife cuts were made using a Micro Feather Ophthalmic Scalpel No. 715 (Electron Microscopy Sciences) which had a triangular blade 7 mm long, 2 mm wide at the base and 0.2 mm thick, with an edge angled 15° to the handle. The knife was mounted on a stereotactic manipulator angled 20° from vertical in the coronal plane, so that the knife edge was angled a total of 35° from vertical. It was positioned at the correct AP coordinate (males: 4.0 mm lateral to the midline on the right side, and then inserted 5.0 mm penetrating at a 20° angle; females: 3.8 mm lateral to the midline on the right side and then inserted 4.75 mm penetrating at a 20° angle). It was then moved 2.5 mm (males) or 2.3 mm (females) medially as measured by the ‘horizontal’ scale of the manipulator, then withdrawn using the ‘vertical’ adjustment of the manipulator. The sham condition followed the same procedure as above except the blade was only inserted 1.0 mm into brain to cut only into the cortex, sparing perforant path fibers.

Immediately following the perforant path fiber transection, two additional craniotomies were drilled over the left and right hippocampi for CTB infusions. Placement of two different Alexa Fluor conjugates of this retrograde tracer (488 and 647) enabled verification of the right hemisphere perforant path fiber transection (488) with intact fiber verification in the left hemisphere (647) within the same animals. Each CTB conjugate was diluted to 1% in PBS (Conte et al., 2009a,b). All CTB infusions were made using a Nanoject II Auto-Nanoliter Injector (Drummond Scientific Company) fit with a glass pipette backfilled with the appropriate CTB conjugate. In male rats the CTB was infused at -3.8 mm posterior to Bregma, ±2.2 mm mediolateral and between -3.4 and -2.1 ventral to the dural surface. The glass pipette was lowered to -3.4 ventral to the dural surface and 50.6 nL of CTB was infused. The pipette was left in place for 1 min and then moved up 100 μm in which another 50.6 nl infusion was placed. This process was repeated at every 100 μm until a final infusion was made at -2.1 mm ventral to dura for a total of 0.708 μL of CTB. This way the tracer infusion included the CA3 and dentate gyrus subregions. Following the final infusion, the pipette was left in place for 150 sec, and then slowly advanced up by 0.5 mm where it was left in place for another 150 sec allowing for the tracer to diffuse away. After the final waiting period, the pipette was slowly removed from the brain. These infusion procedures were then repeated with other CTB conjugate in the contralateral hemisphere. The CTB infusions in the female rats were identical to those used for the males except the coordinates were adjusted to -3.6 mm posterior to bregma, ±2.1 mm mediolateral and -3.3 to -2.0 mm ventral to the dura.

### 2.6 Discrimination Retraining and Rodent Mnemonic Similarity Task Testing

After surgery, rats were given 1 week to recover prior to being retrained to distinguish between two distinct LEGO objects until they reached 81.25% correct. Rats then had 2 days off before the first foil test began. To prevent over-familiarization with the lure objects, rats were tested every 3 days, with 2-3 days off between testing sessions. Rats completed 50 discrimination trials on test days, with 10 having a frog as a control, three sets of 10 trials with objects built out of LEGO blocks with increasing similarity to the target, and 10 trials with two identical copies of the target object as control to ensure that rats were not using any latent cures to detect the correct object. The Lure 1 object had 50% overlapping visible features, Lure 2 had 70% overlap, and Lure 3 had 90% overlap. Performance on this version of the rodent mnemonic similarity task confirmed a behavioral deficit due to the right perforant path transection and these data published previously (Burke et al., 2018).

### 2.7 Compartment Analysis of Temporal Activity with Fluorescence In situ Hybridization (catFISH)

On the final day of behavioral testing rats performed two 5 min behavioral epochs of LEGO block object discrimination, separated by 20 min in which they were returned to the home cage. During one epoch, rats performed the 70% overlap discrimination and in the other epoch rats were tested on the 90% overlap condition, with the order of the different discrimination problems counterbalanced between animals. Due to the two-epoch design on the experiment day, the 50% overlap lure (Lure 1) was not included. See **Figure 1B** for an image of the different stimuli that were used in these two overlap conditions.

Immediately following completion of the second behavioral epoch, rats were placed into a glass bell jar saturated with isoflurane vapor and were rapidly anesthetized prior to being decapitated with a rodent guillotine. Immediately following decapitation, rat brains were extracted, and flash frozen in chilled isopentane (−50°C). Brains were then stored in a freezer at -80°C until cryosectioning. Twenty-micron thick coronal sections of both hemispheres were cut on a cryostat and arranged on Superfrost Plus slides (Fisher Scientific; Hampton, NH) with whole brains from three experimental animals that always included at least one transection and one sham animal. Additionally, a hemisected cage control and a MECS (maximal electroconvulsive shock administered prior to sacrifice as a positive control) animal were included to ensure that slight deviations in staining between slides would not confound the data. The MECS condition served as a positive control to ensure that the *in situ* hybridization worked. This was verified by qualitative examination, and these data were not quantitively analyzed. Tissue was then stored at -80°C until it could be processed using fluorescence *in situ* hybridization (FISH). Every 10^th^ slide was stained with the nuclear counterstain DAPI to examine for projection neurons in the lateral entorhinal cortex. This procedure has been confirmed to be a valid method by which to disconnect the perforant pathway as published previously in Burke et al., 2018.

Twenty-four hours before FISH occurred, brain tissue was moved to a -20°C freezer to prepare for thawing the subsequent day. FISH was performed as previously described (Guzowski et al., 1999; Burke et al., 2005, 2012). Briefly, a commercial transcription kit and RNA labeling mix (Ambion REF #: 11277073910, Lot #: 10030660; Austin, TX) was used to generate a digoxigenin-labeled riboprobe with a plasmid template containing a 3.0 kb *Arc* cDNA (generously provided by Dr. A Vazdarjanova; Augusta University). Tissue was incubated with the probe overnight and *Arc* positive cells were detected with anti-digoxigenin-HRP conjugate (Roche Applied Science Ref #: 11207733910, Lot #: 14983100; Penzberg, Germany). Cyanine-3 (Cy3 Direct FISH; PerkinElmer Life Sciences, Waltham, MA) was used to visualize labeled cells and nuclei were counterstained with DAPI (ThermoFisher Scientific).

### 2.8 Imaging and Analysis

Optical z-stacks were acquired with a Keyence BZ-X7000 all-in-one fluorescence microscope and acquired with the Keyence software (Keyence; Osaka, Osaka Prefecture, Japan). Slides were loaded into the Keyence and were clasped in place. The brain sections were then initially viewed under 2X zoom to verify anatomical location of regions of interest. For the lateral entorhinal cortex, the rhinal sulcus was used a landmark and images were acquired from ventral to the sulcus once the prominent fan cells of layer II were visible. These clearly distinguished entorhinal cortex from area 35 of the perirhinal cortex (Burwell, 2001). Lateral entorhinal cortical Images were acquired from approximately -4.8 to -6.0mm posterior to bregma. For CA3, images were acquired from dorsal CA3 between approximately -3.5 and - 4.2mm posterior to bregma. The Keyence objective was then switched to 40X zoom to acquire optical z-stacks at increments of 1μm. The cell bodies were first viewed under the DAPI channel and exposure was adjusted so that neurons and glia could be differentiated. Additionally, the *Arc* channel was turned on and exposure was adjusted to minimize background while retaining cytoplasmic labeling. For both the cell body (DAPI) and *Arc* channels (Cy3) channels exposure, gain and offset was kept consistent across images within a single slide. Images were taken in both the left and right hemispheres in the areas of interest from 2-3 slides for each animal. For several rats only 2 slides had usable tissue that was not damaged. In the hippocampus, two images were taken from each slide for distal CA3 (area within the hilus of the dentate gyrus), and proximal CA3 (closer to CA2), so that the two images did not overlap. In the lateral entorhinal cortex, a dorsal image (closer to the rhinal sulcus) and a more ventral image were taken. **Figure 2** shows the anatomical regions that images were acquired from.

**Figure 2.**
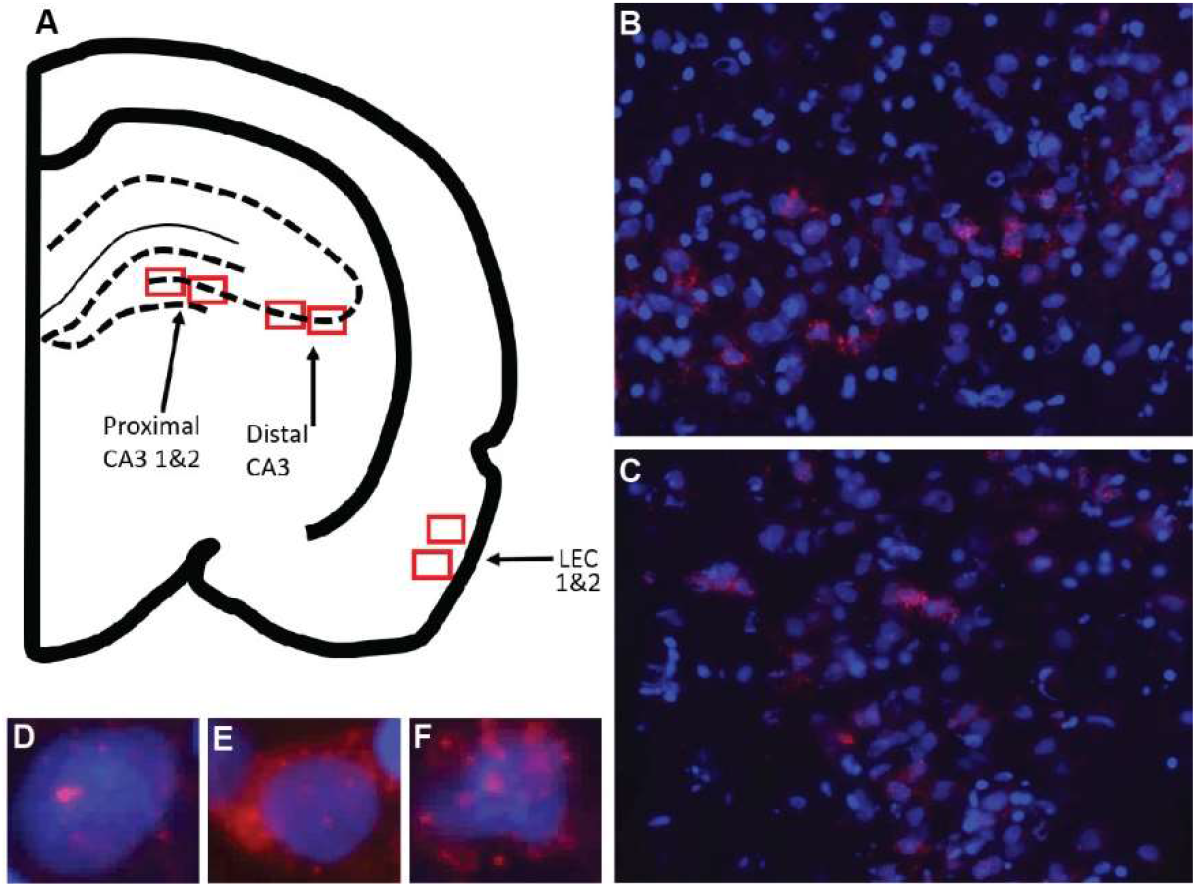
Images taken for analysis of immediate early gene *Arc* expression in CA3 (proximal, distal) and LEC. (**A**) Schematic of the locations for image collection. Two areas were sampled from each region of interest shown by the red boxes. (**B**) Example image of CA3 and (**C**) LEC analyzed for *Arc* expression. Overall cell count was determined, and cells were then classified as being positive for *Arc* in the nucleus or foci (**D**), cytoplasm (**E**), or both foci and cytoplasm (**F**).

Prior to counting, image file names were coded so that experimenters were blind to condition. Initially, the *Arc* channel was turned off to prevent bias while identifying nuclei. Only neurons that were completely contained within the image stack were counted. These neurons were identified by only counting nuclei that were visible in the Median 20% of the Z-stack. Any neurons that were cut off by the edges were also excluded. The *Arc* channel was then turned on and cells were classified as being positive for *Arc* in the nucleus (foci+), in the cytosol (cyto+), in both the nucleus and cytosol (both), or no expression (neg). In order to be considered “foci+”, foci visible in at least 4 adjacent planes. A “cyto+” cell was defined as a cell with *Arc* following the outer contours of at least 30% of the nucleus. A “both” cell was defined as a cell that met the criteria for foci+ and cyto+. A “neg” cell was defined as a cell that did not meet any of these criteria.

### 2.9 Statistical Analyses

Analyses were performed with R Studio and SPSS for Windows. Behavioral and *Arc* expression variables were tested for statistical significance with repeated measures ANOVAs with similarity of object pairs and brain hemisphere as the within-subjects factors and PPT or sham surgery and sex as the between-subjects factors. Potential relationships between behavior and *Arc* expression variable were explored with principal components analysis.

## 3 RESULTS

### 3.1 Mnemonic Similarity Discrimination Testing

Rats performed two behavioral epochs of LEGO block object discrimination testing, counterbalanced between 70% and 90% visual overlap target-lure LEGO object pairs. **Figure 1C** shows the mean percent correct for the Sham and perforant path transection (Cut) groups in the different overlap conditions. Mean performance values for each trial type were entered into a repeated measures ANOVA with lure similarity (70% versus 90%) as a within-subjects factor and sex and experimental surgical group (Sham or Cut) as between-subjects factors. There was a main effect of lure similarity (F_[1,28]_ = 62.361, p < 0.001), and overall rats averaged 90.9% (±1.41 SEM) correct on the 70% similarity trials and 65.2% (±2.83 SEM) correct on 90% similarity trials. Furthermore, there was a main effect of experimental surgical condition (F_[1,28]_ = 6.072, p = 0.020), with Sham animals having more correct trials compared to the Cut rats. Performance did not vary by sex (F_[1,28]_ = 0.752, p = 0.393).

### 3.2 Arc Expression in Relation to Behavioral Epochs

Animals that remained in their home cage for the duration of the experiment and performed no behavioral assessments showed low *Arc* expression (Total Activity in the LEC: 5.81%; CA3 distal: 3.87%; CA3 proximal: 4.17%). **Figures 3A-D** show the average population activity between the different overlap epochs inferred from *Arc* expression in proximal and distal regions of CA3 in both the left (Intact) and right (Cut) hemispheres. A repeated measures ANOVA revealed no main effect of Object Similarity (F_[1,55]_ = 0.3435, p = 0.560), Surgical Condition (F_[1,55]_ = 0.5277, p = 0.471), Hemisphere (F_[1,55]_ = 0.0252, p = 0.875), or Sex (F_[1,55]_ = 1.5889, p = 0.213) on CA3 *Arc* expression. Furthermore, there were no interaction effects between any of the between- or within-subjects factors (p > 0.364 for all comparisons).

**Figure 3.**
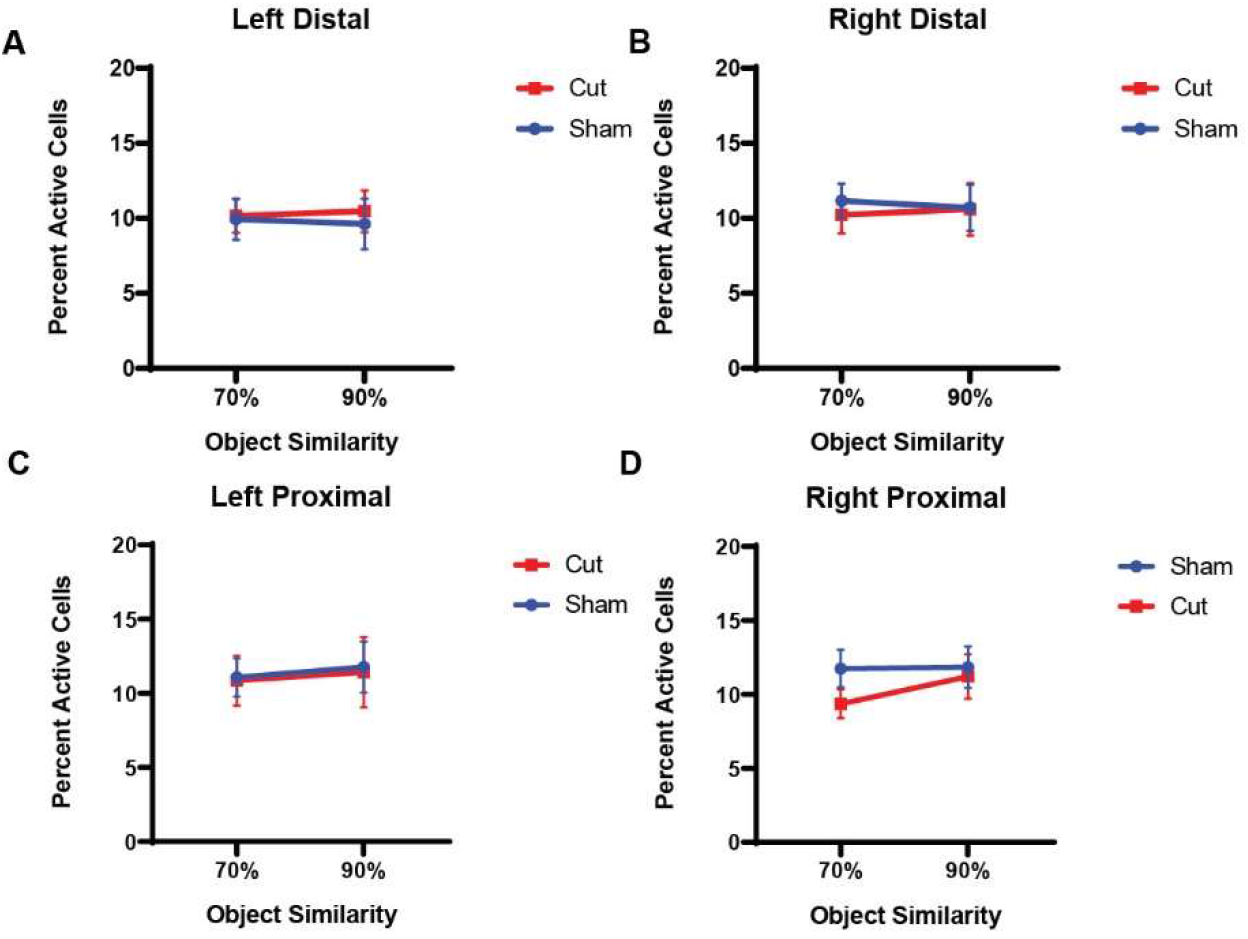
Percentage of active cells, as determined by expression of the immediate early gene *Arc*, in the CA3 subregion of the hippocampus. There were no within- or between-subjects effects found for either distal (**A, B**) or proximal (**C, D**) CA3.

Figure 4. shows the average population activity in the left (A) and right (B) hemispheres of the LEC. A repeated measures ANOVA showed a main effect of the within-subjects factor of Object Similarity (F_[1,55]_ = 5.8482, p = 0.019) and between-subjects factor of Hemisphere (F_[1,55]_ = 4.28338, p = 0.044), but no effect of surgical Condition (Sham versus Cut: F_[1,55]_ = 0.04222, p = 0.838) or Sex (F_[1,55]_ = 0.00322, p = 0.955). The significant between-subjects effect of hemisphere indicated that overall population activity was higher in the left hemisphere of the LEC in all animals compared to the right. This observation of activity differences between the left and right LEC suggests that the cortical damage induced by the surgical procedures in both the Sham and Cut conditions led to reduced LEC activity. Furthermore, in both surgical groups, *Arc* expression during the 90% overlap discrimination condition was lower compared to 70% feature overlap condition (F_[1,55]_ = 5.8482, p = 0.019). This could be due to the more unique features of the LEGO objects in the 70% condition resulting in more neuron activation, in contrast to the 90% condition in which the two objects are more likely to be perceived as identical.

### 3.4 Similarity Scores

In order to investigate the degree to which population activity overlapped across the 70% and 90% overlap behavioral epochs, while accounting for differences in the size of neuronal ensembles that were positive for *Arc* mRNA, similarity scores were calculated. The similarity scores between epochs 1 and 2 were calculated as:

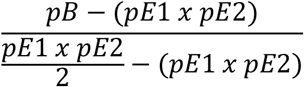

where pB represents the proportion of cells active during both epochs, pE1 the proportion of cells active during epoch 1, and pE2 those active during epoch 2. **Figures 5A-D** show the similarity scores across proximal and distal CA3 in both the left and right hemispheres. A repeated measures ANOVA revealed that males had significantly higher similarity scores than females (F_[1,27]_ = 6.1465 p = 0.020). Furthermore, there was not a significant interaction between condition and hemisphere (F_[1,27]_ = 2.926, p = 0.099), but the interaction between sex and surgical condition did reach statistical significance (F_[1,27]_ = 4.5518, p = 0.042). The reasons for these sex differences will require additional exploration. Figure 5E shows the similarity score in the left and right hemispheres of the LEC across surgical conditions. A repeated measures ANOVA found no main effect of or interaction effects between the factors of Hemisphere, Condition, or Sex on LEC similarity scores (p ≥ 0.351 for all comparisons).

**Figure 4.**
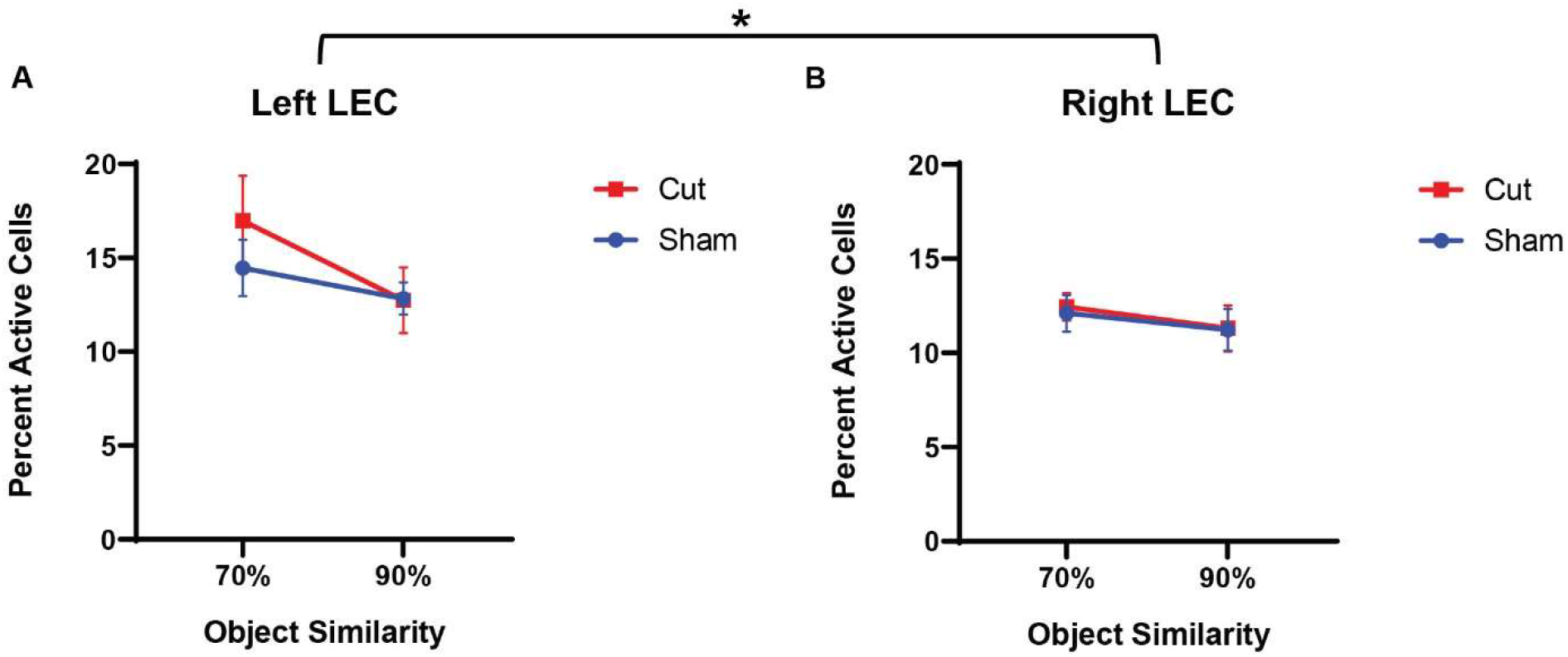
Percentage of active cells, as determined by expression of the immediate early gene *Arc*, in the Left (**A**) and Right (**B**) Lateral Entorhinal Cortex. Quantified *Arc* expression in the left LEC was higher than the right hemisphere (F_[1,51]_ = 4.283, p = 0.044), but there was no effect of surgical condition or sex. Activity was lower in both hemispheres and across surgical conditions for 90% similarity trials compared to 70% similarity (F[1,55] = 5.8482, p = 0.019).

**Figure 5.**
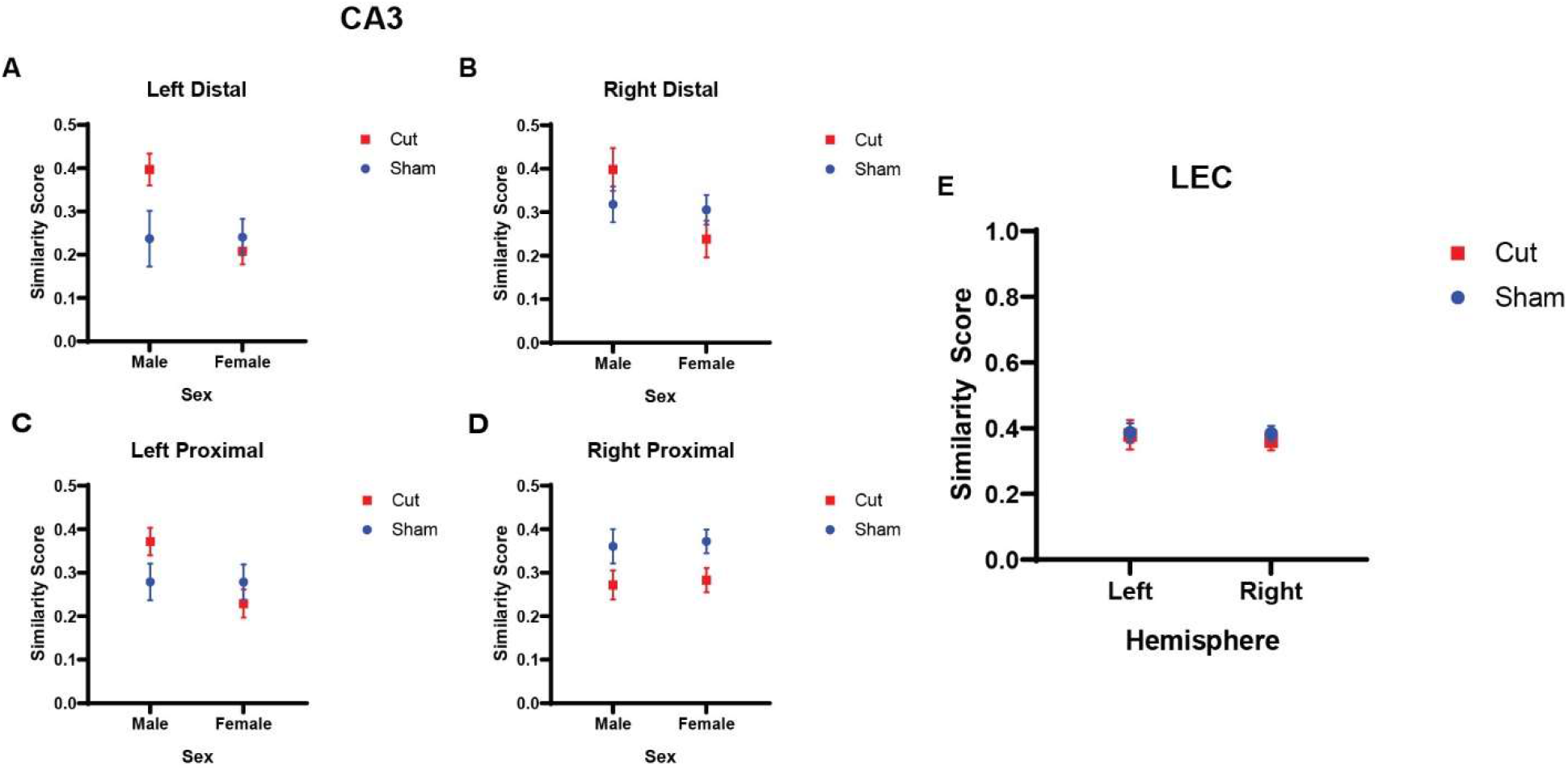
Similarity score of population activity between epochs 1 and 2 as evaluated by *Arc* expression. (**A-D**) CA3 similarity scores did not show a main effect of hemisphere. There was a main effect of sex (F_[1,27]_ = 6.1465 p = 0.020), and an interaction effect between sex and surgical condition (F_[1,27]_ = 4.5518, p = 0.042). (**E**) LEC similarity scores showed no main effect of hemisphere (F[_1,25]_ = 0.903, p = 0.351), sex (F[_1,25]_ = 0.139, p = 0.712), or surgical condition (F[_1,51]_ = 0.057, p = 0.813).

### 3.5. Relationship between Arc Expression Patterns and Behavioral Performance

To interrogate whether there was a potential relationship between the expression patterns of the immediate early gene *Arc* in any of the hippocampus regions of interest and discrimination performance, data reduction was performed with a Principal component analysis (PCA) that included all measures of *Arc* expression. A PCA with varimax rotation was performed due to split loadings; however, one variable was still shown to have cross-loadings for components 1 and 3, thus was dropped and the PCA was run again. According to the total variance explained (see Table 1), the model using the top 6 components (eigenvalues > 1.0) explained 77.613% of variance in the data. The first component (eigenvalue = 3.670) corresponded to activity during the 70% object similarity trials and accounted for 21.590% of variance in the data. Increased *Arc* transcription in both left and right hemisphere distal and proximal CA3 regions positively loaded onto the first component (≥ 0.660), and similarity score for the right LEC negatively loaded onto component 1 (−0.606). The second component (eigenvalue = 2.777) corresponded to activity during the 90% object similarity trials and uniquely accounted for 16.333% of variance in the data. Increased *Arc* transcription in both left and right hemisphere for distal CA3 as well as right proximal CA3 regions positively loaded onto the second component (≥ 0.584). The third component (eigenvalue = 2.215) corresponded to left hemisphere CA3 similarity scores and accounted for 13.027% of variance. Higher similarity scores (values approaching 1) for both proximal and distal CA3 subregions positively loaded onto component 3 (≥ 0.793). *Arc* expression in the LEC during both overlap conditions loaded positively (≥0.730) onto the 4^th^ component and accounted for 11.021% of the variance. CA3 similarity score negatively loaded onto the component (−0.644). The variables that loaded onto the 5^th^ and 6^th^ components did not have an obvious relationship to each other and accounted for less than 10% of the variance.

**Table 1.** Rotated Component Matrix with Varimax rotation from Principal Component Analysis.

| **Principal Component Analysis** |  |  |  |  |  |  |
| --- | --- | --- | --- | --- | --- | --- |
| **Component** | **1** | **2** | **3** | **4** | **5** | **6** |
| Eigenvalue;<br>% Variance | 3.670;<br>21.590% | 2.777;<br>16.333% | 2.215;<br>13.027% | 1.874;<br>11.021% | 1.477;<br>8.689% | 1.182;<br>6.953% |
| Left Proximal CA3 70% | 0.851 |  |  |  |  |  |
| Right Distal CA3 70% | 0.747 |  |  |  |  |  |
| Left Distal CA3 70% | 0.718 |  |  |  |  |  |
| Right Proximal CA3 70% | 0.660 |  |  |  |  |  |
| Right LEC Similarity Score | -0.606 |  |  |  |  |  |
| Right Proximal CA3 90% |  | 0.874 |  |  |  |  |
| Right Distal CA3 90% |  | 0.858 |  |  |  |  |
| Left Distal CA3 90% |  | 0.584 |  |  |  |  |
| Left Distal CA3 Similarity Score |  |  | 0.917 |  |  |  |
| Left Proximal CA3 Similarity Score |  |  | 0.793 |  |  |  |
| Left LEC 70% |  |  |  | 0.794 |  |  |
| Left LEC 90% |  |  |  | 0.730 |  |  |
| Right Distal CA3 Similarity Score |  |  |  | -0.644 |  |  |
| Right Proximal CA3 Similarity Score |  |  |  |  | -0.875 |  |
| Left LEC Similarity Score |  |  |  |  |  | -0.788 |
| Right LEC 90% |  |  |  |  | 0.533 | 0.547 |
| Right LEC 70% |  |  |  |  |  |  |

To evaluate any relation between *Arc* activity and behavioral performance, a two-tailed Pearson’s Correlation score was calculated between the regression factors for components 1 and 2 and the corresponding behavioral epoch performance (**Table 2, Figure 6**). The test between the first factor, corresponding with *Arc* activity during 70% object similarity trials, and animals’ percent correct trials during the 70% overlap epoch did not show a correlation between *Arc* expression and performance (Pearson’s r = 0.141, p = 0.512). Interestingly, the same analysis of the second regression factor from the PCA, which corresponded to activity during the 90% object similarity trials, and animals’ percent correct trials during the 90% overlap epoch revealed a significant negative correlation between *Arc* expression and discrimination performance (Pearson’s r = -0.463, p = 0.023), indicating that animals with more CA3 activity had worse performance on the 90% feature overlap condition of the mnemonic similarity task.

**Table 2.** Pearson’s Correlations between percent of correct trials during 70% or 90% overlap conditions and either (**A**) The Regression factor score for PCA Component 1 or (**B**) Regression factor score for PCA Component 2. * Indicates significant effect.

| **Pearson's Correlations** |  |  |  |  |
| --- | --- | --- | --- | --- |
| **A** |  | **Percent Correct**<br>**70% Similarity** | **Percent Correct**<br>**90% Similarity** | **PCA Component**<br>**1** |
| **Percent Correct**<br>**70% Similarity** | Pearson<br>Correlation | 1 | 0.000 | 0.141 |
|  | Significance (2-tailed) |  | 0.999 | 0.521 |
|  | N | 32 | 32 | 24 |
| **Percent Correct**<br>**90% Similarity** | Pearson<br>Correlation | 0.000 | 1 | 0.053 |
|  | Significance (2-tailed) | 0.999 |  | 0.805 |
|  | N | 32 | 32 | 24 |
| **PCA Component**<br>**1** | Pearson<br>Correlation | 0.141 | 0.053 | 1 |
|  | Significance (2-tailed) | 0.512 | 0.805 |  |
|  | N | 24 | 24 | 24 |
| **B** |  | **Percent Correct**<br>**70% Similarity** | **Percent Correct**<br>**90% Similarity** | **PCA Component**<br>**2** |
| **Percent Correct**<br>**70% Similarity** | Pearson<br>Correlation | 1 | 0.000 | 0.067 |
|  | Significance (2-tailed) |  | 0.999 | 0.757 |
|  | N | 32 | 32 | 24 |
| **Percent Correct**<br>**90% Similarity** | Pearson<br>Correlation | 0.000 | 1 | -0.463* |
|  | Significance (2-tailed) | 0.999 |  | 0.023 |
|  | N | 32 | 32 | 24 |
| **PCA Component**<br>**2** | Pearson<br>Correlation | 0.067 | -0.463* | 1 |
|  | Significance (2-tailed) | 0.757 | 0.023 |  |
|  | N | 24 | 24 | 24 |

**Figure 6.**
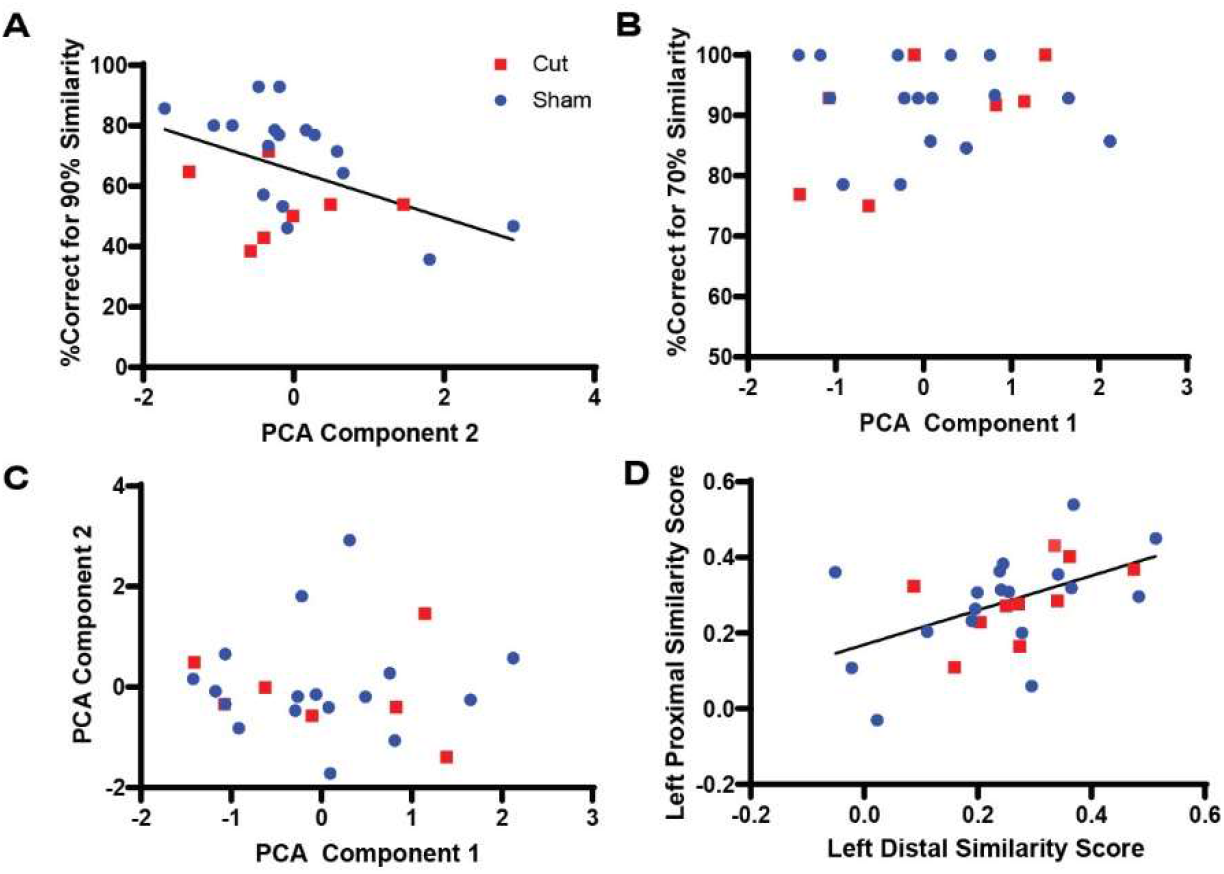
Correlations between regression scores from PCA analysis and corresponding behavioral epoch performance. (**A**) Comparison of PCA Components 1 and 2, which correspond to Arc expression during the 70% and 90% similarity trials, respectively (Pearson’s r = 0.0, p = 1.0). (**B**) Comparison of PCA Component 1 and percent correct trials during 70% similarity trials yielded no significant correlation (Pearson’s r = 0.141, p = 0.512). (**C**) Comparison of PCA Component 2 and percent correct trials during 90% similarity trials showed a significant correlation (Pearson’s r = -0.463, p = 0.023). (**D**) Comparison of left distal and proximal CA3 similarity scores showed a significant correlation (Pearson’s r = 0.512, p = 0.004).

## 4 DISCUSSION

This study tested the hypothesis that perforant path fiber damage leads to increased neuronal activity in CA3, which has been reported in aged animals (Wilson et al., 2005; Yassa et al., 2011b; Robitsek et al., 2015; Thomé et al., 2016; Lee et al., 2021a) and older adults (Yassa et al., 2011b; Yassa et al., 2011a; Reagh et al., 2018). A previous study reported that unilateral damage to the perforant path impaired the ability to discriminate between a target and 90% similar lure (Burke et al., 2018). During the experimental procedures of the current study, all animals were able to correctly discriminate between a target object and 70% similar lure object significantly more than when the lure had 90% feature overlap with the target object. Moreover, there was a significant difference in performance accuracy based upon surgical condition, with animals that received perforant path transection making more errors than the animals that underwent sham surgery (**Figure 1C**). While the fiber transection resulted in behavioral impairments, there were no significant effects of surgical condition on *Arc* expression in CA3 or the LEC. Additionally, there were no sex differences in behavioral performance or percent of active cells during either the 70% or 90% overlap conditions. Together these observations suggest that a unilateral loss of perforant fibers does not significantly alter neuronal activation in CA3 and LEC, despite this surgical treatment leading to altered behavioral performance. These observations lend further support to notion that intrinsic circuits of the hippocampus are reliant on local computations and are therefore robust against alterations in afferent input (Brun et al., 2008; Zutshi et al., 2022).

When the data from both surgical groups were analyzed together, *Arc* activation was related to participation in the behavioral task. Specifically, in both hemispheres of the LEC there was more *Arc* expression associated with the 70% overlap condition of the mnemonic similarity task compared to the 90% overlap condition. There are at least two potential explanations for this observation. First, rats received more rewards in the 70% overlap condition compared to when the feature overlap was 90%. It is conceivable to more rewards resulted in more LEC activation. In support of this idea, the LEC receives direct innervation from dopamine neurons in the ventral tegmental area (Loughlin and Fallon, 1984; Oades and Halliday, 1987; Björklund and Dunnett, 2007). Furthermore, a recent study reported that neurons in the LEC were active during the acquisition of cue-reward association (Lee et al., 2021b). An alternative, but not mutually exclusive possibility is that in the 70% overlap condition, the two objects are more likely to be perceived as distinct, while the 90% overlap objects are more likely to be perceived as identical. By this account, two unique objects could produce more LEC activation then 2 identical objects, which have fewer unique features. Previous studies have reported that LEC neurons recorded from rats are responsive to objects (Deshmukh and Knierim, 2008, 2011; Tsao et al., 2013; Knierim et al., 2014), which supports the notion that when an animal experiences more unique objects, a larger proportion of LEC neurons will be active. In contrast to the LEC, the proportion of neurons in CA3 that were positive for *Arc* mRNA did not differ between the 70% and 90% feature overlap conditions.

Similarity score was used to quantify the extent that transected versus sham animals showed differences in the population overlap of active neuronal ensembles between the different overlap conditions. This measure accounts for differences in overall activation and the expected overlap based on chance (Vazdarjanova and Guzowski, 2004). Scores range from 0 to 1, with a higher similarity score indicating greater population overlap in cells active between behavioral epochs. For the LEC, the similarity of population activity between the 70% and 90% overlap condition was around 0.4 for all conditions and did not differ by hemisphere, sex, or surgical condition. For CA3, there were differences in the similarity score by surgical condition that interacted with sex. For left proximal, left distal and right distal subregions of CA3, similarity score was higher for male compared to female rats, but this effect was only evident in the rats with right perforant path transection. The reason for the sex difference is not apparent and requires further investigation, but it could be due to the transection affecting males more than females, which has been reported for ventral hippocampal lesions (Beninger et al., 2009). In the right proximal CA3 subregion, the similarity score was lower in the rats that received perforant path transection. As this area of CA3 does not receive direct entorhinal cortical input (Witter et al., 2000), the reason for this difference is not evident.

Dimension reduction was performed to examine what *Arc* expression variables were related to each other. The proportion of neurons expressing *Arc* during the 70% overlap condition in all subregions of CA3 positively loaded onto first principal component. This is consistent with the anatomy of CA3 that contains strong autoassociative projections (Amaral and Witter, 1995; Witter, 2007), which could support similar activity patterns among different neurons in the CA3 circuit. The first principal component did not correlate with performance during the 70% overlap condition, which could be due to the near ceiling level of performance during this condition that did not allow for sufficient parametric space to detect neural-behavioral relationships. The second principal component was positively associated with the proportion of CA3 neurons that expressed *Arc* during the 90% overlap condition in right distal and proximal CA3, as well as left distal CA3. Importantly, the second principal component positively correlated with performance on the 90% overlap condition of the mnemonic similarity task. This observation suggests that more activity in CA3 supports the ability to discriminate between stimuli that share features. Importantly, total activation in CA3 was between 8-12% for all animals. If these data are considered within the context of previous research showing worse performance on the mnemonic similarity task with increased CA3 activity (Yassa et al., 2010; Maurer et al., 2017), it suggests that there is an inverted U-shaped relationship between CA3 activity and the ability to discriminate between stimuli that share features. Discrimination accuracy will wane if CA3 activity is too low or two high (Kent et al., 2016).

Notably the present data call into question current models of pattern separation and completion (Marr, 1971; Rolls, 2013). These network models assume that patterns of brain activity represent unique states and that the hippocampus generates sharp transitions in activity between two well-learned states as input gradually changes (McNaughton and Morris, 1987; Guzowski et al., 2004; Leutgeb et al., 2005). According to the pattern separation and pattern completion theory, the hippocampus makes binary decisions, but would be incapable of reconciling situations in which small changes must be tracked and incorporated within a continuous context (Maurer and Nadel, 2021). To solve the mnemonic similarity task, animals need to compare a current stimulus to a learned target to determine if it is identical, while simultaneously detecting small differences in the configuration of features in order to detect and ignore the foil object. This task cannot be adequately solved with a single binary pattern completion or separation computation, but rather is more likely to be supported by re-entrant loops, and amorphous networks that can monitor stimulus equivalence while tracking perceptual continuities and discontinuities (Maurer and Nadel, 2021). Furthermore, the current experiment challenges the theory of pattern separation and completion as we did not observe large differences in orthogonalization of population activity overlap between the LEC and CA3. When population overlap between epochs was quantified, the similarity scores for neuronal ensembles in LEC and CA3 were comparable. While it is conceivable that the dentate gyrus would have shown reduced overlap, it is unclear as to why input patterns were be orthogonalized by the dentate gyrus to then become more similar in CA3.

Overall, this project shows that a unilateral transection of the perforant path does not recapitulate age-related changes in neural activity as measured by immediate early gene *Arc* expression. This suggests that while perforant path degradation is a feature of aging and may contribute to age-related cognitive decline, it does not seem to be the catalyst for age-related changes in CA3 activity patterns. These results therefore beg the question of whether perforant path dysfunction is more easily compensated for in younger animals or if other factors common to advanced age compound with perforant path degradation in a unique way to cause cognitive deficits seen in older populations. To further untangle the processes involved in neural compensation and cognitive resilience, it will be important to examine if there are effects of a perforant path transection on other regions known to be involved in object recognition such as the perirhinal cortex or CA1 region of the hippocampus. If these regions are found to have higher levels of activation as measured by *Arc* expression, it could indicate a compensatory effect that mitigates any disruption of cognitive function due to loss of perforant path fibers from the lateral entorhinal cortex.

## Acknowledgements

Funding provided by Evelyn F. McKnight Brain Research Foundation, NIH/NIA research grant R01AG055544, and the T32AG020499-16A1. We acknowledge Michael G. Burke for maze construction, and Kim Robertson and Sarah Lovett for administrative help.

